# Borealization: a biodiversity refugium and evolutionary cradle during the Permian–Triassic climate crisis

**DOI:** 10.64898/2026.10.12.758598

**Authors:** Adam Shatwell, William J. Foster

## Abstract

Understanding how organisms track suitable environments during rapid climate change is constrained by the brevity of modern observations. Using the fossil record, we identify the mid- to high-latitude Boreal Ocean as a regional biodiversity refugium during the Permian–Triassic climate crisis. Additional Boreal successions indicate sustained diversity and comparatively hospitable conditions. Biogeographic reconstructions reveal that 30% of Sassendalen taxa survived the extinction event, with two-thirds expanding or shifting their latitudinal ranges into the Boreal Ocean. More than 90% of newly evolved species originated there, and their lineages subsequently dispersed at differing rates into other regions throughout the Early Triassic. These findings identify poleward migration and high-latitude refugia as key drivers of biodiversity persistence and evolutionary recovery during climate crises.

## Main Text

Understanding the factors that govern ecosystem resilience to environmental disturbance is central to predicting/forecasting and mitigating the impacts of ongoing climate change (*1*). Among the most important are the breadth of an organism’s environmental tolerances, their geographic distribution, and their capacity to shift ranges to track their preferred niche (*2*). Poleward range shifts and expansions have been documented in the warming modern oceans using ecological time-series (*3, 4*), but long-term inferences are impeded by short (annual to decadal-scale) focal durations. Consequently, predictive ecosystem models are largely calibrated against modern and historical datasets, inheriting this temporal constraint (*5*). The sedimentary record extends these observations across centennial to millennial scales and captures ecosystem responses to other climate crises. These records provide natural experiments for identifying mechanisms that promote biodiversity persistence and recovery under extreme environmental change, thereby informing predictions of future biotic responses (*1, 6*). Here, we reconstruct the biogeographic origins and distributions of fossil taxa recovered from Sassendalen, Svalbard, to test whether the Boreal Ocean functioned as a regional biodiversity refugium and evolutionary cradle during and following the Permian-Triassic climate crisis, enabling the persistence of complex marine ecosystems.

The Permian-Triassic climate crisis, 251.9 Ma, was an extreme high magnitude, high-rate warming event eradicating an estimated 81% of marine species within a maximum duration of 60,000 years, making it the most severe mass extinction (*7–8*). The crisis is widely attributed to the emplacement of basaltic sills from the Siberian Traps Large Igneous Province into carbon-rich sediments, increasing atmospheric CO_2_ concentrations six-fold (*9-10*). Marine extinctions are hypothesized to have resulted from the combined effects of the ‘deadly trio’ of thermal stress (>10°C equatorial sea-surface warming), widespread ocean deoxygenation and ocean acidification, with greenhouse conditions persisting throughout the Early Triassic (*11-12*). Recovery of the global biosphere was similarly protracted, particularly for benthic ecosystems, requiring several million years (*13*).

Most Permian-Triassic transition fossil assemblages are known from numerous low-paleolatitude sections within the former Tethys Ocean (Fig. 1A). Griesbachian (earliest Triassic) Tethyan communities are species-poor, dominated by a few cosmopolitan generalists, and lack multiple trophic levels, extensive bioturbation and metazoan reef ecosystems (*13*). More complex ecosystems are not known until the succeeding Dienerian, Smithian and Spathian substages, more than one million years post-extinction (*14-15*). However, many genera and higher clades are Lazarus taxa. Accounting for Lazarus taxa, only one ecological mode of life likely perished across the mass extinction despite the depauperate state of known earliest Triassic communities (*16*). This ‘hidden complexity’ has been variously attributed to scarcity of Lower Triassic taphonomic windows (e.g. silicified faunas) (*17*), survival of populations sizes too small to be preserved (18), or persistence within environmentally favorable refugia until global conditions improved (*19*). Distinguishing among these hypotheses has remained challenging.

**Fig. 1.**
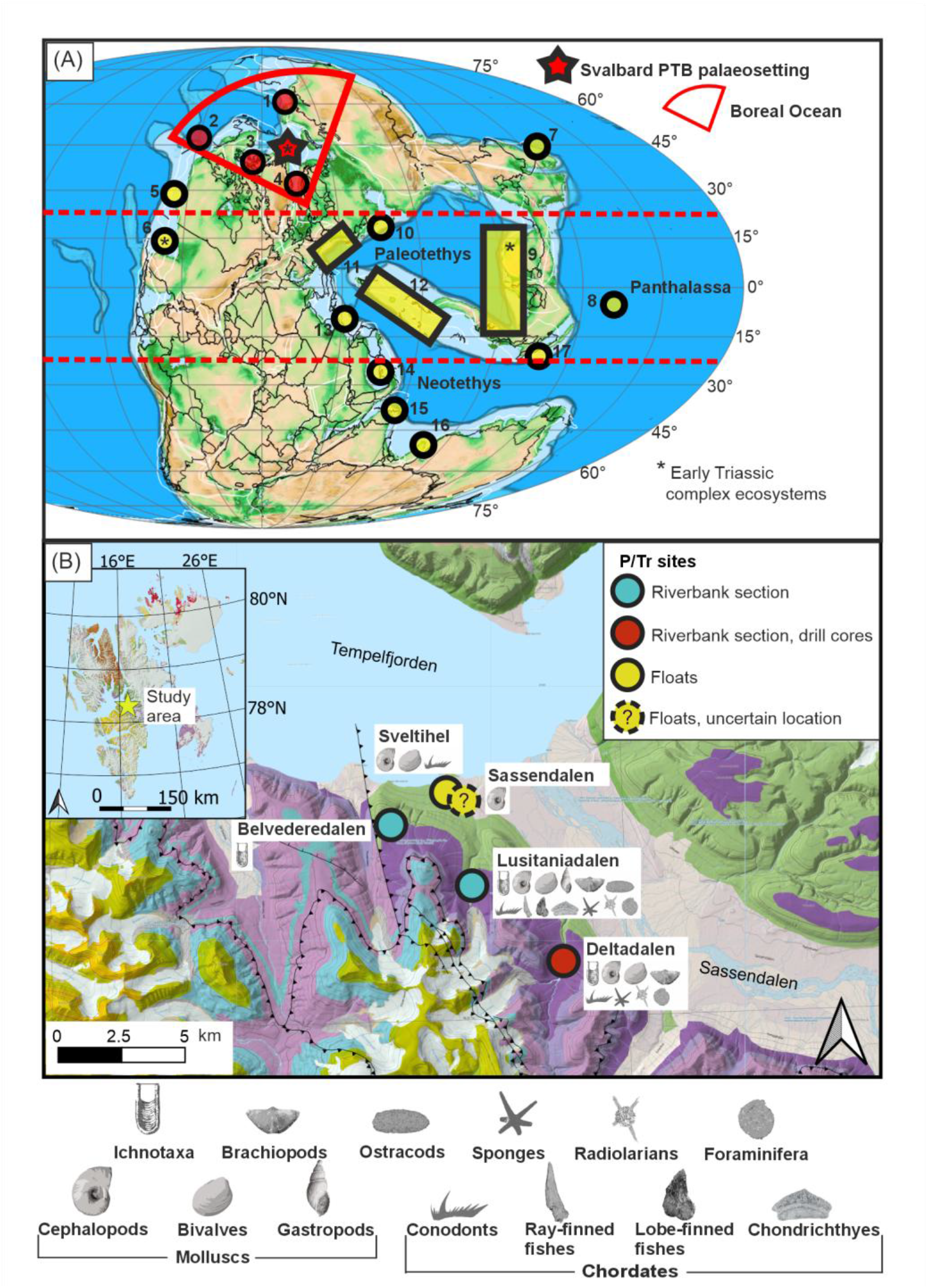
(A) Global palaeogeographic distribution of the main study areas for Early Triassic marine ecosystems, represented by numbered circles (red = within Boreal Ocean). Rectangles mark broader continuous regions of interest. Asterisks show locations of known complex ecosystems from the Early Triassic (Induan and Olenekian ages). Areas are numbered as follows: Boreal Ocean - 1) Verkhoyansk; 2) northeastern Canada (Sverdrup Basin); 3) eastern Greenland; Panthalassa – 4) western Canada; 5) western USA, with Paris Biota; 6) south Primorye; 7) Japan (ophiolite terranes); Tethys – 8) South China Block and Indochina, including Guiyang Biota; 9) Caucasus; 10) Alps, Carpathians and Dinarides (northern Italy, Slovenia, Hungary, Croatia, Bosnia and Herzegovina, Serbia); 11) Cimmeria (Iran, Armenia, Azerbaijan); 12) Taurides (southern Turkiye); 13) Oman; 14) Pakistan, northern India, Kashmir, Tibet; 15) western Australia; 16) Sukhothai (Malaysia). PaleoDEM from (61), map exported from GPlates v2.5.0 (62). **(B)** Locations and fossil assemblages of each of the PTB-spanning sections, drill cores and floats within the Sassendalen area. Green and purple areas represent Permian and Triassic outcrop extents. Key of fossil groups at base of figure. (Inset) Geological map of the Svalbard archipelago, the gold star marking the study area. Basemaps from the Norwegian Polar Institute (WGS 84/Arctic Polar Stereographic), assembled within QGIS (v3.34.12).

Here we find that the relatively isolated Boreal Ocean supported complex and diverse marine ecosystems even throughout the climate crisis. The latest Changhsingian – early Griesbachian Sassendalen Biota comprises over 40 families spanning seven phyla. Biogeographic reconstructions indicate that many surviving taxa were new arrivals or ‘warm guests’ from lower paleolatitudes, whereas newly-evolved species were largely Boreal Ocean endemics. Geochemical proxy records consistently indicate comparatively habitable environmental conditions. Together, these findings identify the Boreal Ocean as both a regional high-latitude biodiversity refugium and an evolutionary cradle in the aftermath of the climate crisis. We term this higher-latitude persistence and reassembly of complex ecosystems Borealization, reflecting its prior use to describe poleward migration of modern marine communities (*4*).

### Svalbard and the Boreal Ocean

The Boreal Ocean formed a northwestern embayment of Panthalassa along the margin of Pangaea and was geographically and climatically distinct from the low-latitude Tethys Ocean. Fossiliferous shelf deposits outcrop across modern Alaska, northern Canada, eastern Greenland, northern Russia and Svalbard, but remain far less studied than their coeval Tethyan or tropical Panthalassan successions (*20*). In central Svalbard, laterally extensive siliciclastic exposures of the Permian-Triassic Vikinghøgda Formation preserve an exceptional record of ecosystem recovery following the Permian-Triassic mass extinction (*21–23*). Its lowermost unit, the offshore Deltadalen member, was deposited at 50-55°N (*24*). A pronounced negative δ^13^C excursion, accompanied by the abrupt demise of trace fossils, bioturbation, and chert, alongside the proliferation of extinction lipid biomarkers, which marks the Permian-Triassic mass extinction and climate crisis approximately corresponds to the contact between the Vikinghøgda and Kapp Starostin Fm. (*25-27*). The Permian/Triassic boundary occurs 2.8 m higher, as defined by the first appearance of the conodont *Hindeodus parvus* (*28*).

### An earliest-Triassic complex ecosystem

The Sassendalen biota represents the oldest known complex marine ecosystem following the Permian-Triassic mass extinction, and the only such ecosystem documented from the immediate aftermath of the extinction, i.e., occurring precisely at the Permian/Triassic boundary. To characterize its taxonomic and ecological structure, we compiled 354 body-fossils and 404 ichnofossil occurrences from the lowest 15 m of the Deltadalen Mb. and uppermost 6 m down into the Kapp Starostin Fm. Most occurrences derive from the riverbank sections at Lusitaniadalen, Deltadalen and Belvederedalen, supplemented by two drill cores (*28*) and stratigraphically constrained float material (*29-30*) (Fig. 1B).

In total, we identified 116 species, including 66 formally named species, representing 48 genera, 45 families, 15 classes and seven phyla, together with 16 ichnogenera. All but two species occur exclusively above the mass extinction horizon. The assemblage comprises sponge spicules, radiolarians, conodont elements, vertebrate teeth and bones, brachiopods, ostracods and ammonoids together with silicified bivalves, gastropods and agglutinated foraminifera (Fig. 2). Most fossils are concentrated within calcareous concretions, whereas ichnofossils and foraminifera are known throughout the mudstones. Species richness substantially exceeds that of the more intensely sampled Griesbachian assemblages from the Great Bank of Guizhou (South China) and the Werfen Formation (southern Alps) (*31*). This exceptional diversity reflects the co-occurrence of benthic, planktonic and nektonic clades, a feature rarely documented in earliest Triassic ecosystems (*32*). Furthermore, phylum-level richness matches, and class-level and lower taxonomic richness exceeds, that of the Early Triassic Guiyang and Paris biotas, the only other described mixed-community Lagerstätten from low paleolatitudes (*14-15*).

**Fig. 2.**
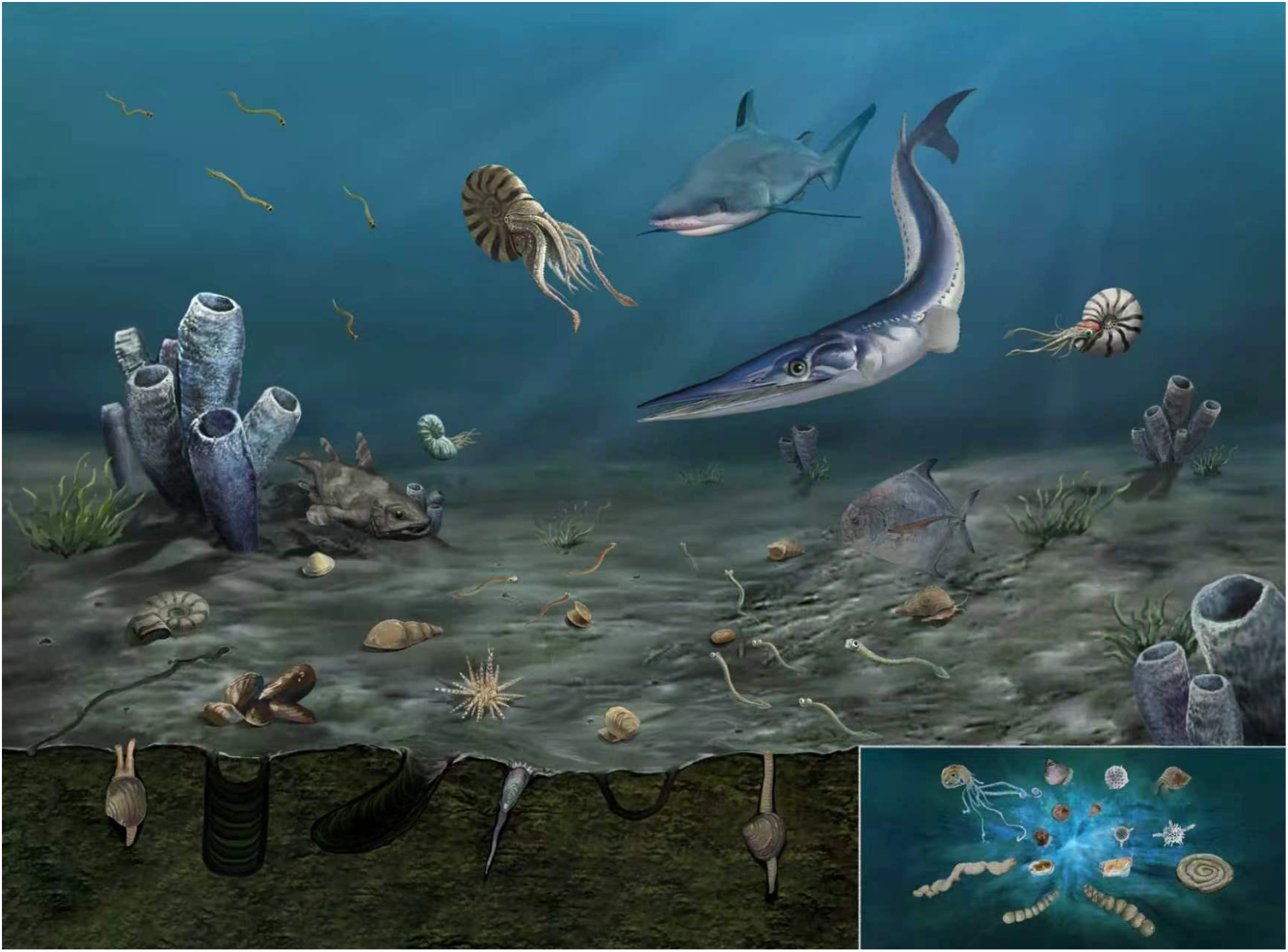
Artistic reconstruction of the Sassendalen Biota and insert showing the microfauna, each depicted organism represents an identified species (illustration by Huang Peng and Ji Jiao).

The Sassendalen Biota is constrained to the oldest Triassic magnetostratigraphic and biostratigraphic zones, based on ammonoid, palynomorph and conodont zonations (*24,27, 33*). U-Pb zircon ages from ash beds immediately above the Permian/Triassic boundary at Deltadalen (252.13 ± 0.62 Ma), confirm an earliest Triassic age (*28*). Using the sedimentation rate of 61m/Myr estimated for the Deltadalen Member (*28*), we constrain the Sassendalen Biota to within 240kyrs of the mass extinction horizon, spanning the latest Changhsingian and early Griesbachian. This is likely an overestimate, as the inferred c. 60kyr mass extinction duration could itself cover up to 3.66m of the basal Deltadalen Mb. at the inferred sedimentation rate, and possible seasonal varves observed in the DD-1 core indicate intervals of substantially more rapid sedimentation. The Sassendalen Biota therefore predates all previously described post-extinction complex marine ecosystems by at least ∼0.9 Myr.

Using a relatively coarse exospace scheme, the Sassendalen taxa occupied at least eleven modes of life, ranging from infaunal sediment miners and erect sessile suspension feeders to free-swimming pelagic carnivores. Predators included large ammonoids (up to 40 cm diameter) and the large predatory fishes *Saurichthys, Bobasatrania* (Actinopterygii) and *Palaeobates* (Hybodontiformes). Although these genera are known elsewhere in the Early Triassic, Sassendalen preserves their oldest post-extinction occurrences (*34*) and isolated fish teeth extend their record below the Permian/Triassic boundary (*35*). Bone remains of an identified coelacanthid have also been observed from the Sassendalen Biota, representing the only known occurrence of this drift-hunting ecology globally at this time, but also coelacanths are reported as reaching 2-3 m body size later in the Triassic in Svalbard (*36*). These observations indicate that complex food webs, including meter-scale secondary consumers with high metabolic rates, remained present during the climate crisis. High primary productivity is independently supported by lithology-independent enrichment of eukaryotic phytoplankton biomarkers (*27*). Ichnofossil diversity likewise increases up-section, documenting recolonization of deep infaunal (>5cm) habitats (*22*), a pattern also recognized elsewhere in the Boreal Ocean (*20, 37*). Ecological specialization is further indicated by the encrusting brachiopod *Orbiculoidea*, the durophagous *Bobasatrania* and *Palaeobates*, and two commensal symbiotic agglutinated foraminiferal genera associated with hexactinellid sponges. *Reophax spiculifer* tests are exclusively constructed from spicules, while *Tolypammina vagans* is interpreted as an endoparasite because it is commonly preserved within sponge tissues (*38*). The persistence of these specialized interactions implies sustained populations of biomineralizing taxa despite strong extinction-driven selection against marine calcifiers and silicifiers.

Although the Sassendalen biota has a diagenetically silicified component, this taphonomic window alone cannot account for its exceptional ecological complexity or that of the wider Boreal Ocean. Multiple modes of preservation, including original silica, phosphate, and calcite recrystallisation, together with the ichnofossil assemblage, independently document tiered benthic communities comprising detritivores, suspension feeders, predators and plankton irrespective of silicified fossils. Silicification likely enhanced the recorded diversity of benthic mollusks, as observed in the late Griesbachian Wadi Wasit Block fauna (*39*), and of agglutinated foraminifera, but cannot explain the presence of diverse trophic guilds or specialized ecological interactions. Likewise, the occurrence of large predators and other specialists argues against their persistence at globally negligible population sizes that precluded reliable fossilization (*18*). Instead, the exceptional complexity of the Sassendalen Biota is most parsimoniously explained by genuine regional differences in ecosystem recovery, consistent with the development of a Boreal refugium.

### A Boreal refugium

A key characteristic of refugia is that they experience reduced exposure to the dominant drivers of environmental disruption, allowing persistence of immigrant taxa and facilitating regional ecosystem recovery. In the Boreal Ocean, the abrupt loss of bioturbators across the mass extinction horizon followed by rapid Griesbachian recolonization in both shoreface and offshore transition deposits indicates environment degradation that was substantial but comparatively limited (*20, 40*). Among the Permian-Triassic ‘deadly trio’ of extinction mechanisms, preserved Sassendalen fossils show no evidence of acidification-related dissolution or etching (*41*). Although climate models predict amplified warming at higher latitudes, absolute temperatures remained lower than those in tropical regions (*42*). Redox-sensitive metal enrichments and pristane/phytane ratios from Lusitaniadalen, used as indicators of seafloor oxygenation, record only transient dysoxia and euxinic conditions near the extinction interval followed by rapid recovery (*26, 40*). These patterns agree with proxy records from shallower western Svalbard sections (*37*) (fig. 3). Enhanced oxygenation may have resulted from frequent storm-driven mixing (*20*), with prolonged oxygen stress potentially restricted to deeper offshore environments.

**Fig. 3.**
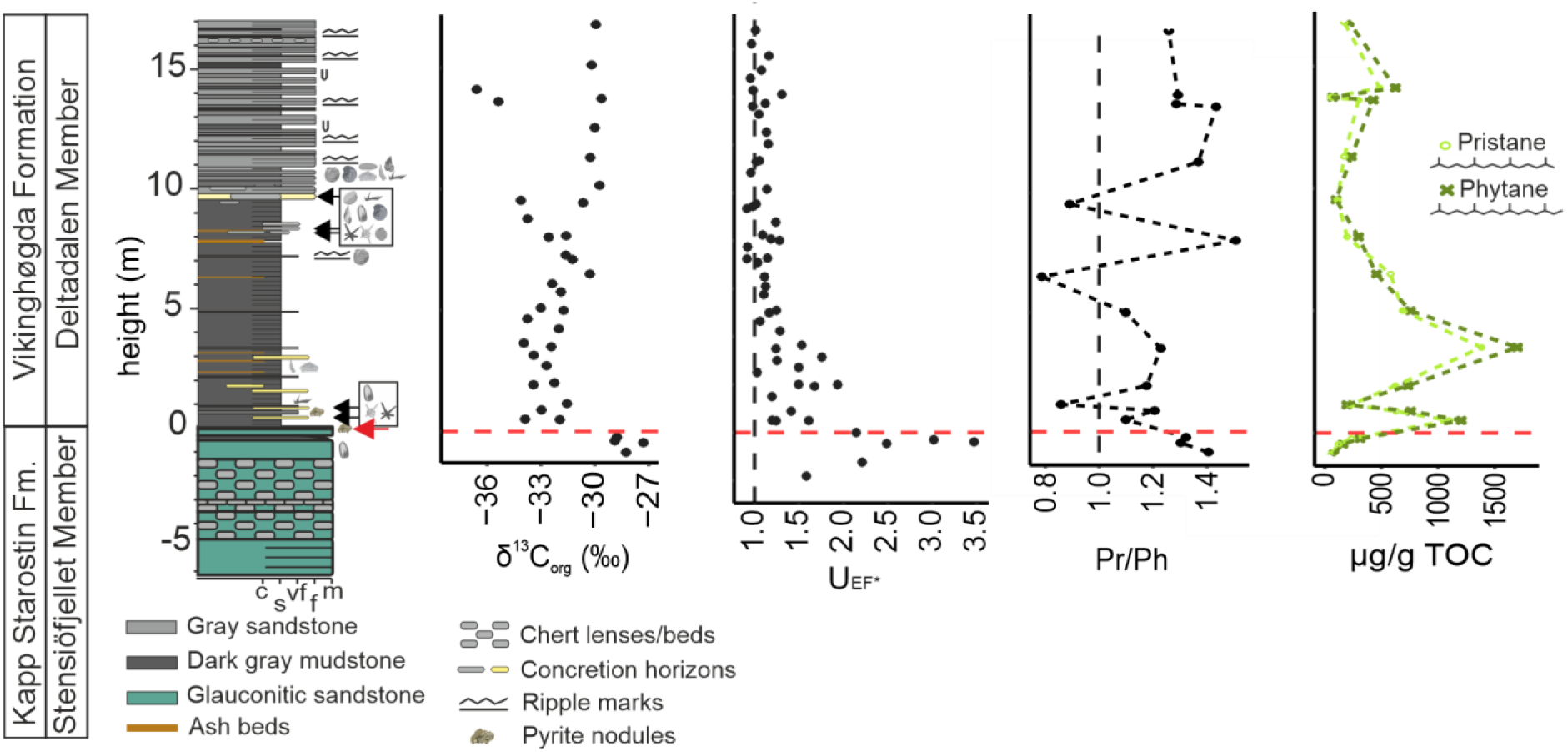
Organic geochemical proxy records from Lusitaniadalen, δ^13^C_org_, U_EF*_, pristane/phytane, and abundance of pristane and phytane (µg/g TOC). U_EF_ indicates redox conditions, Pr/Ph values greater than 1 indicate oxic conditions, and increased pristane and ohytane abundance indicate increased primary productivity. Data after Buchwald et al. (40).

A key expectation of refugia is the presence of eponymous refugees: taxa that occupied more environmentally stressed regions before the disruption and subsequently tracked suitable conditions through range expansion, migration, or contraction. To determine whether such climate migrants occurred in the Sassendalen Biota, we compiled all global occurrences of the 64 named species recovered from the basal Deltadalen Mb. and assessed their stratigraphic relationship to the Sassendalen Biota (Fig. S1). We classified these taxa as either mass extinction survivors or post-extinction radiations, and further categorized survivors according to their geographic distributions and range changes across the crisis (Table S2).

Overall, 30% of the Sassendalen species survived the major climate perturbation (fig. 4). Nine survivors had no Boreal Ocean record: some remained present elsewhere during the interval and represent range expanders, whereas others shifted their distributions into the Boreal Ocean and represent range shifters. Of the ten survivors with pre-extinction Boreal records, three had cosmopolitan distributions before the crisis that subsequently contracted to the Boreal Ocean. Therefore, 63% of survivor taxa underwent biogeographic changes associated with entry into, or restriction to, the Boreal Ocean. These migrants span multiple groups, including agglutinated foraminifera, radiolarians, bivalves, ostracods and conodonts. Only one Sassendalen species, *Orbiculoidea winsnesi*, was endemic to the Boreal Ocean across the extinction event. Although extensive silicification in the uppermost Kapp Starostin Fm. limits quantitative assessment of local extinction patterns, diverse Permian assemblages of brachiopods, bryozoans, ostracods, conodonts and sponges occur within scattered calcareous horizons and the siliciclastic succession below (*43*). None of these taxa are recorded from the Sassendalen Biota.

**Fig. 4.**
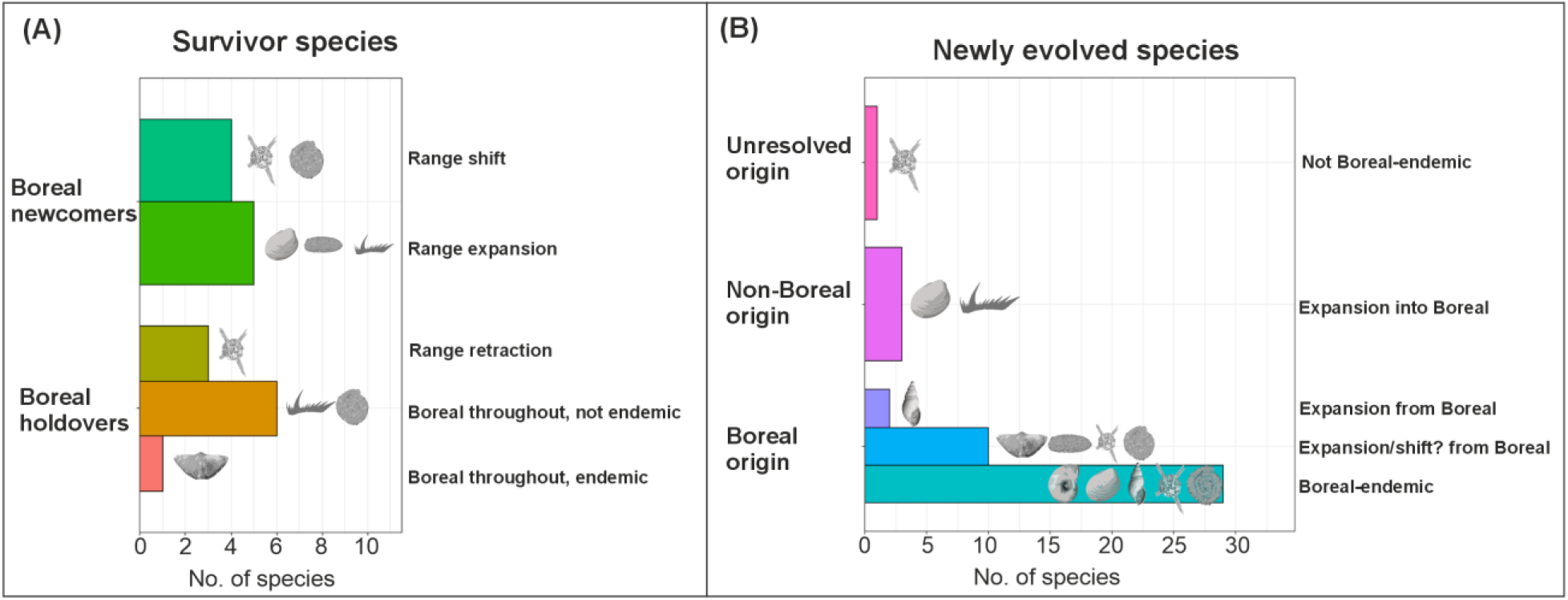
Sassendalen taxon richness within the (A) ‘survivor’ and (B) ‘newly evolved’ categories and their subdivisions. Icons (key in figure 1) show which major fossil groups are represented within each subdivision. Sponges, ray-finned fishes and cartilaginous fishes excluded due to lack of species-level identification.

Sassendalen range-expanders were exclusively metazoans, including pelagic conodonts and the eurytopic bivalve *Claraia*. Although nektonic taxa can be highly sensitive to climate-related stressors, their mobility and homeostatic capability may have facilitated persistence through the Permian-Triassic crisis by enabling redistribution away from deteriorating environments and access to transient refugia (*19, 44*). In contrast, range-shifting species comprised only foraminifera and radiolarians, while all three range-retractions were radiolarians. This distinction may reflect narrower thermal tolerances among rhizarian lineages relative to metazoans a pattern previously linked to extinction severity across the Permian-Triassic transition (*44-45*).

Radiolarians appear particularly sensitive to warming and oxygen limitation, potentially explaining the displacement or regional disappearance of formerly widespread species (*44-45*). Their strong biogeographic response is consistent with elevated extinction rates and pronounced provincialism among the limited Early Triassic faunas (*46*). In contrast, Sassendalen foraminifera were dominated by hypoxia-tolerant agglutinated clades, which may have enabled persistence and migration across broader shelf environments, explaining both Boreal survivorship and absence of range-contractions among these taxa (*47*).

Boreal Ocean survivors without range contraction were predominantly cosmopolitan taxa and included nektonic conodonts, hypoxia-tolerant foraminifera and eurytopic generalists *Claraia* and *Orbiculoidea*. The abundance of these widespread taxa, together with numerous immigrants, coupled with geochemical evidence for comparatively hospitable condition and the near-absence of pre-extinction endemics, indicates that environmental disruption remained severe despite regional buffering. Extinction rates within the Boreal Ocean were therefore likely substantial, consistent with predictions based on clade-specific thermal and oxygen tolerances (*42-45*).

However, diversity losses appear to have been largely offset by poleward niche-tracking of tropical taxa, resulting in high community turnover while maintaining unexpectedly diverse and ecologically complex ecosystems.

### A diversity cradle

To assess whether the Boreal Ocean also acted as a center of post-extinction diversification, we reconstructed the geographic origins of newly evolved Sassendalen species. Of the 45 named newly evolved species, 91% first appear in the Sassendalen Biota or other Boreal successions, indicating that they originated within the Boreal Ocean (Fig. 4). These include all newly evolved foraminifera and nine of ten newly evolved radiolarian species, and all mollusks except 3 widespread bivalves. Furthermore, 71% of newly evolved species remained endemic to the Boreal Ocean throughout their tenure, including 21 of the 25 named mollusk species and all ten ammonoid species. Such pronounced species-level endemism is unexpected given the weak global provincialism of the earliest Triassic (*46*) and suggests that comparatively hospitable conditions promoted *in situ* diversification while limiting dispersal until environmental conditions improved elsewhere. Two Boreal gastropod species subsequently expanded into the Paleotethys, whereas most endemic lineages remained regionally restricted. As these endemics account for much of the elevated species richness of the Sassendalen Biota, the Boreal Ocean appears to have functioned as an earliest Triassic evolutionary cradle. This pattern contrasts with the conventional ‘out-of-the-tropics’ model and instead suggests that extreme extinction severity and persistent environmental stress temporarily shifted the center of marine diversification to higher latitudes (*48*). Conodonts provide a notable exception, as there relatively low extinction rates may explain why Sassendalen conodont and ammonoid assemblages are dominated by range expanding survivors, whereas co-occurring ammonoids comprise exclusively newly-evolved endemics – despite broadly similar nektonic lifestyles (*49*).

Direct survival and post-extinction diversification meant that 66% of named genera within the Sassendalen Biota persisted across the climate crisis. By sustaining both survivor and newly radiating lineages, the Boreal Ocean provided refuge for several clades that subsequently shaped Mesozoic marine ecosystems. Although some Paleozoic groups, such as the bellerophontid gastropods, disappeared within the following five million years (*50*), others, including architectibranch gastropods, diversified and persisted to the present day. Two bivalve families evolved in the Boreal Ocean, and one order – Solemyida – is a Griesbachian Lazarus clade when excluding boreal occurrences (*51*). Ceratitid ammonoids, absent from the Boreal Ocean during the Late Permian, radiated there following the extinction before dispersing into lower latitudes from the late Griesbachian, with xenodiscoids giving rise to all subsequent Mesozoic ammonoids

(*52*). Similarly, nearly all Griesbachian bony fish material known, nearly half of known chondrichthyan occurrences, and the oldest marine reptiles are concentrated in Boreal successions (*53*). The oldest marine reptile remains are also known from central Svalbard, albeit from a younger deposits (*23*). In contrast, bryozoans, four crinoid orders, three radiolarian families, and microconchid reefs remained largely confined to the Boreal Ocean throughout the Early Triassic (*54-55*). These contrasting dispersal histories suggest that ecological traits, particularly mobility, influenced the rate at which recovering lineages expanded beyond this high-latitude evolutionary center.

### Borealization: a mechanism of hyperthermal recovery

Our findings support a general mechanism of ecosystem recovery following hyperthermal climate crises. Rather than relying solely on *in situ* survival, biodiversity and ecosystem functions can be maintained through the redistribution of taxa into environmentally buffered higher-latitude basins. We term this process Borealization: the concentration of global biodiversity and ecological functions within high-latitude refugia through range shifts, expansions, and contractions, followed by *in situ* diversification and the subsequent dispersal of recovering lineages as environmental conditions improve elsewhere. By integrating biogeographic reconstructions with ecological and geochemical evidence, we show that migration and adaptation acted together to sustain complex marine ecosystems through the most severe mass extinction in Earth’s history.

Borealization also has implications for predicting biological responses to ongoing climate warming. Poleward migration is already restructuring modern marine ecosystems (3, 4), and higher-latitude thermal refugia may similarly buffer biodiversity during future climate change where suitable migration pathways remain available (2). However, our results demonstrate that refugia are dynamic rather than static reservoirs of biodiversity. Despite comparatively hospitable conditions, the Boreal Ocean experienced substantial ecological turnover, with widespread immigrant taxa largely replacing pre-crisis regional endemics. Climate refugia may therefore preserve ecosystem function and evolutionary potential while simultaneously eroding unique regional biodiversity, emphasizing that resilience depends on both environmental buffering and the capacity of species to disperse and adapt.

## Acknowledgments

This work builds on more than a decade of collaborative research. We sincerely thank our scientific colleagues whose contributions to the generation, analysis, and publication of the underlying datasets laid the foundation for the present study. In addition, we are grateful to Jørn Hurum for the unpublished identification of a coelacanth from our material. AS is additionally thankful to Grace Shephard, Valentin Zuchuat, Joseph Botting and Kim Senger for their additional feedback on this project.

## Author contributions

Conceptualization: WJF, AS. Methodology: AS, WJF. Investigation: AS, WJF. Visualization: AS, WJF. Funding acquisition: WJF. Project administration: Supervision: WJF. Writing – original draft: AS, WJF. Writing – review & editing: As, WJF.

## Data, code, and materials availability

All data, code, and materials used in the analysis are either available in the supplement or available via Zenodo

